# Paramecium Learning: New Insights and Modifications

**DOI:** 10.1101/225250

**Authors:** Abolfazl Alipour, Mohammadreza Dorvash, Yasaman Yeganeh, Gholamreza Hatam

## Abstract

Learning is a fundamental process in neural systems. However, microorganisms without a nervous system have been shown to possess learning abilities. Specifically, *Paramecium caudatum* has been previously reported to be able to form associations between lighting conditions and cathodal shocks in its swimming medium. We have replicated previous reports on this phenomenon and tested the predictions of a molecular pathway hypothesis on paramecium learning. Our results indicated that in contrast to the previous reports, paramecium can only associate higher light intensities with cathodal stimulation and it cannot associate lower light intensities with cathodal stimulation. These results found to be in line with the predictions of the previously proposed model for the molecular mechanisms of learning in paramecium which depends on the effects of cathodal shocks on the interplay between Cyclic adenosine monophosphate concentration and phototactic behavior of paramecium.

## Introduction

learning is a fundamental process in neural systems and much effort had been devoted to elucidating its mechanisms. Specifically, learning in unicellular organisms is an intriguing observation that has not been investigated adequately. Examples of learning in unicellular organisms include learning in the giant slime mold *Physarum polycephalum* that can learn to predict subsequent cold shocks after a periodic cold shock stimulation (1) or move to colder areas to find food (2). Additionally, in smaller organisms such as *Escherichia coli*, the organism had been reported to be able to predict subsequent carbon sources through the proper expression of genes (3) and shift from fermentation to respiration in yeast (3).

In particular, *Paramecium caudatum* is another unicellular organism reported to show intelligent behaviors such as Spontaneous alternation behavior which requires remembering the previous choice in a T-maze(4) and learning (5, 6). Altogether, these observations suggest that learning might not be restricted to strengthening/weakening of synaptic connections and “intracellular learning mechanisms” may exist in some organisms.

Therefore, investigation of learning in unicellular organisms might reveal fundamental mechanisms of learning that had been preserved throughout the evolution. Specifically, paramecium is an ideal organism to be used in addressing this issue since there has been a long history of research on paramecium’s learning behavior dating back to 1911 by Day and Bentley (7).

Meanwhile, a consensus on presence of learning in paramecium is still lacking. Different attempts to show learning in paramecium faced counter results by other researchers and findings on this topic are still equivocal. One of the latest reports on paramecium by Armus et al. (6) suggested that paramecium can learn to associate different light intensities in their swimming medium to attractive electrical shocks (8, 9). More specifically, single paramecia were observed while swimming in a trough with two bright and dark sides. The organism would receive attractive cathodal shocks when it entered the bright/dark side of the trough depending on the trial. At the end, paramecium had been reported to remember the side of the trough in which is received the attractive cathodal shocks regardless of it being dark or bright (6).

However, the molecular mechanism for this process is still unknown. We have previously suggested a molecular model (10) to explain this behavior based on molecular pathways that link cAMP concentration to the phototactic behavior of paramecium.

However, our model predicted that paramecium cannot learn to associate lower light intensities (the dark side) with cathodal shocks. As such, the main goal of this study is to first replicate previous findings of Armus et al. and second, to test the predictions of the existing hypothesis and reveal the mechanisms of learning in paramecium.

## Materials and methods

### Culture media

Hay-infusion was used as the culture medium for paramecium. Hay was boiled in purified water for 45 minutes and the resultant extract was used for paramecium culture as described below.

#### Paramecium caudatum

specimens Local samples of the Khoshk River in Shiraz (Iran) have been gathered and incubated in hay-infusion as the nutritious culture medium for paramecia. After 3 days, the specimens were checked for the presence of the Paramecium and have been isolated for further evaluation. *Paramecium caudatum* was identified based on its unique morphological features i.e. the large size (300 micrometers) and presence of only one micronucleus beside the large macronucleus.

**Figure 1.**
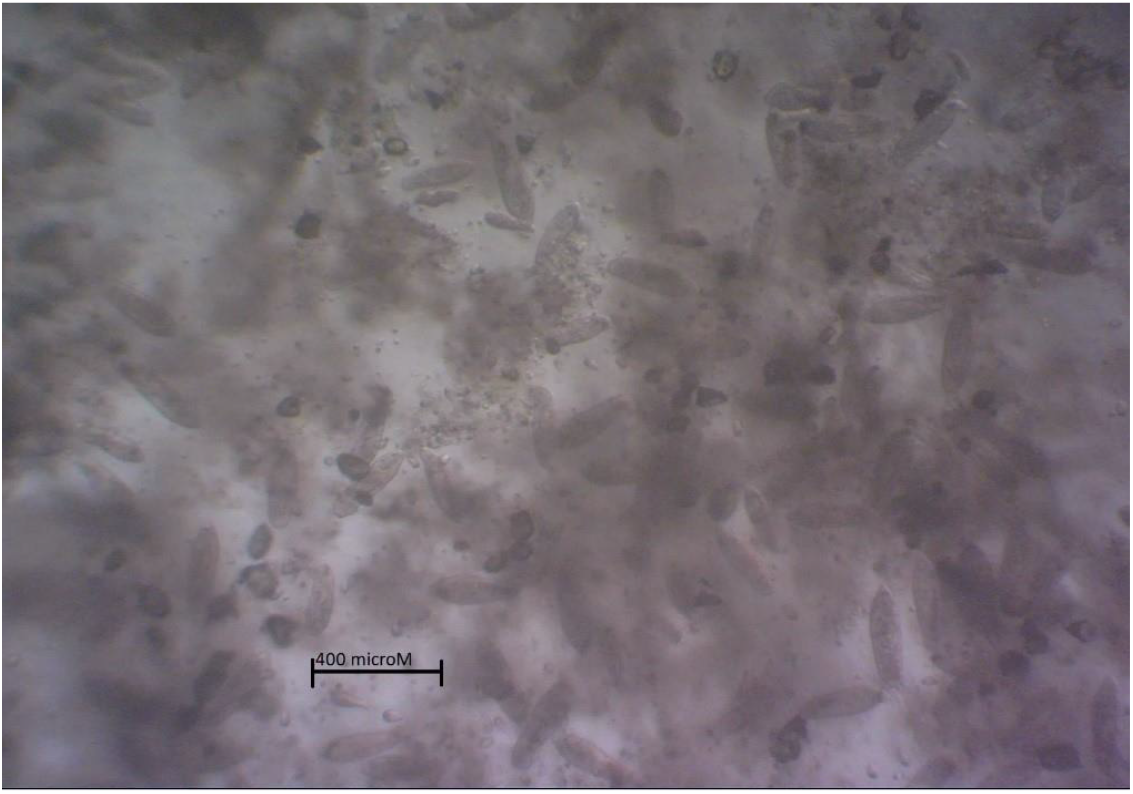
*Paramecium caudatum’s* culture

### Electrical shock device

A microcontroller was used to deliver shocks to the culture medium. (ATMEGA 16 AVR controller). The microcontroller was programmed to deliver 60-millisecond shocks with 500-millisecond no-shock intervals. This circuit was used to deliver cathodal shocks (5 volts, 1 milliamp) to the culture medium.

### Paramecium learning experiment

The methodology of Armus et al was used to investigate the learning behavior of *P. caudatum*. A U-shaped plastic trough (20mm length, 5mm width, and 5mm depth) was filled with the filtered culture medium through a 0.22-micrometer filter. The trough was divided into two dark and light sides using a dark transparent sheet placed under the trough. Light intensity was set to 805±30 and 335±30 candelas for the bright and dark side of the trough, respectively. 84 Paramecia (*P. caudatum)* was divided into three groups of “light association” (n=23), “dark association” (n=26), and “control” (n=36).

For the experiment, each paramecium underwent ten 90-second trials, 7 training trials and 3 test trials for all groups. In training trials of the light association group, each paramecium received electrical shock only when it was on the bright side of the trough. In training trials of the dark association group, each paramecium received electrical shock only when it was on the dark side of the trough. Members of the control group did not receive any shock on either side of the trough. In the test trials, paramecia did not receive any shocks in any of the groups. Additionally, the total time that paramecium spent in the light and dark sides of the trough was recorded for all of the groups.

**Figure 2.**
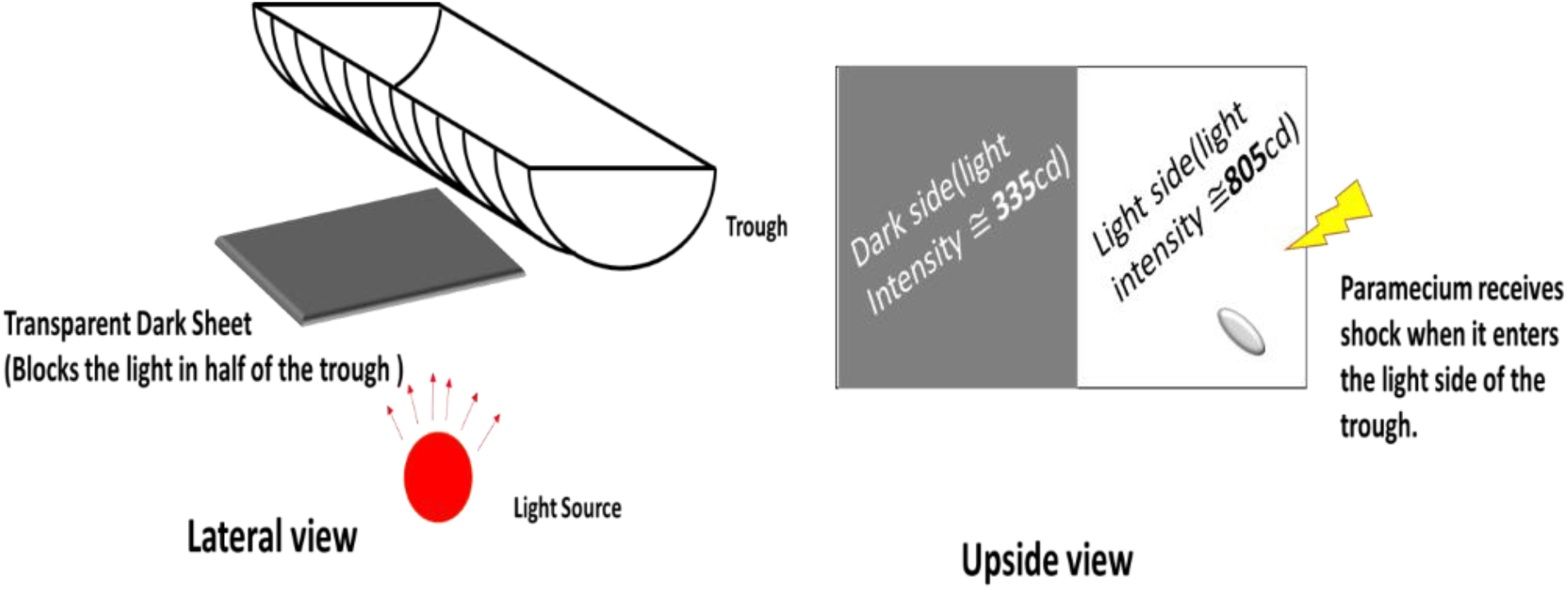
A schematic representation of the experimental setup

## Results

### Light association group

The total duration of the time spent on the light side of the trough was 152.7±12.8 and 105.3± 8.1 seconds for experimental and control group, respectively. The independent *t*-test showed a significant difference between the time spent on the light side of the trough by experimental group comparing with the control group (p<0.01). See Figure 3 for more detail.

**Figure 3.**
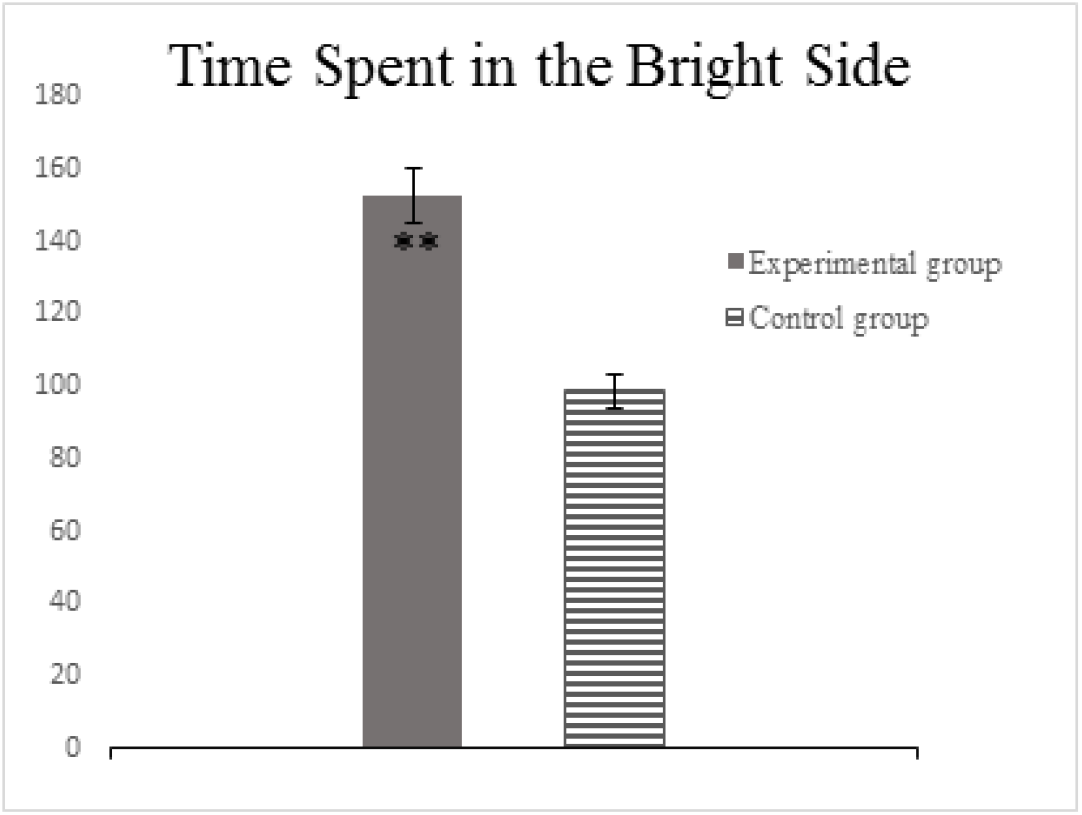
Comparison between the total time spent in the bright-cathodal half of the trough in test trials between control and light association groups. There was a significant difference between two groups (Paired *t*-test, p<0.01).

### Dark association group

The total duration of the time spent on the dark side of the trough was 175.2± 11.7 and 164.6± 8.1 seconds for experimental and control group, respectively. The independent *t*-test showed that the difference between the time spent on the dark side of the trough by experimental group comparing with the control group is not significantly different (p>0.05). See Figure 4 for more detail.

**Figure 4.**
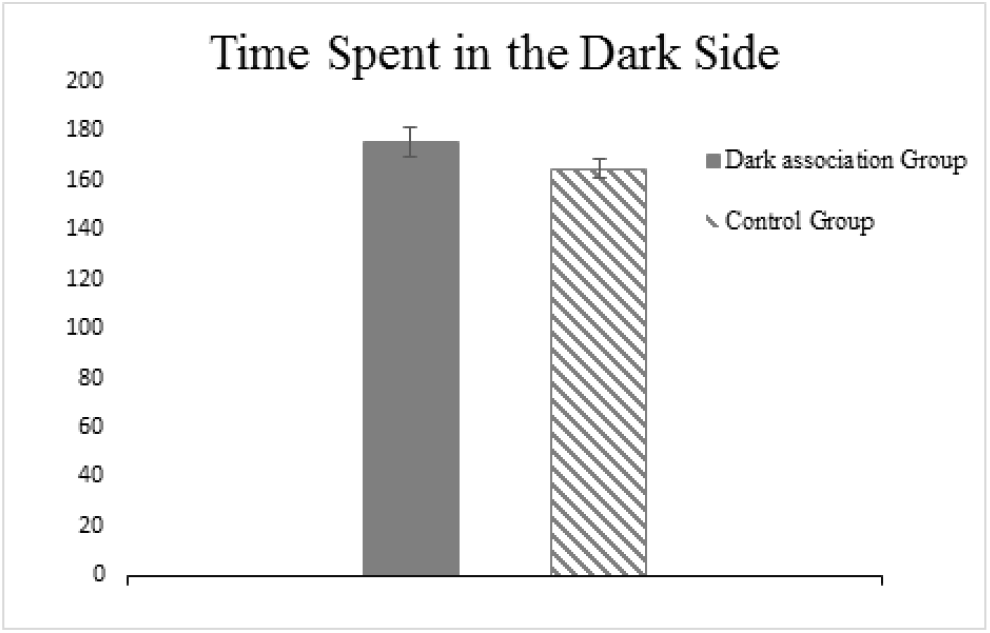
Comparison between the total time spent in the dark-cathodal half of the trough in test trials between control and dark association groups. Two groups did not show a significant difference (Paired *t*-test, *p*>0.05).

## Discussion and conclusion

The present study confirms the existence of learning capabilities in *P. caudatum*. Our study only replicated the core finding of Armus et al. and not all of the reported observations. In the same vein, there are some key points that need to be properly addressed before drawing a conclusion on learning in paramecium. The most important point is that in Armus et al. report(6), there was not a distinction between the paramecia who supposedly learned to associate “dark side” with the cathodal shock and “light side” with cathodal shock. The relationship between light and dark with cathodal shock was simply counterbalanced in their study. Accordingly, the present study made a distinction between dark-cathode and light-cathode association and found that the learning happens only in light-cathode conditions. The theoretical aspects of this issue will be addressed further in this section (using our previous hypothesis (10) on learning mechanisms of paramecium. Particularly, it seems that the data for the control group in Armus et al. report might be unreliable (see Figure 5). Based on Armus et al. report, the control group spent approximately 30 seconds of a 90-second trial in “cathodal” side of the trough. However, since the time spent in the cathodal half is time spent on the dark side of it for 50% of the time and light side for the other 50% of the time, paramecia in the control group should spend on average “45 seconds” of their time in the cathodal side, not 30. Interestingly, the difference between experimental and control group lies within this 15 second time window (6).

**Figure 5.**
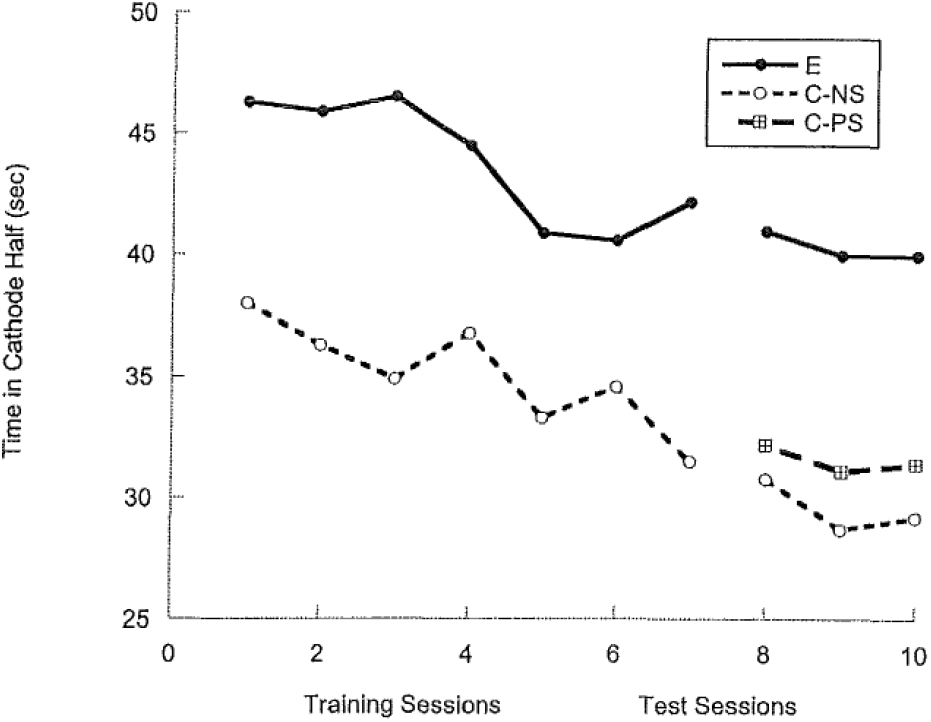
Data from Armus et al. [6]. Each data point indicates time spent in “cathodal half” of the trough in different trials for different groups. The difference between control group and experiment group lies within a 15-second time interval (the test sessions’ data). However, since the data for the control group is the average of time spent in both dark and bright halves of the trough, it should equal to a number close to 45± a possible SD. Taking this consideration into account, the control group’s data seems to be unreliable. Adapted with permission from [6].

Moreover, there are some additional factors that should be considered. First, paramecium spends a significantly longer time in the cathodal half of the trough but this can happen due to an accumulation of unknown substances at the tip of the cathode electrode. To examine this possibility, Armus et al. ran a second control group in which the paramecium was constantly stimulated regardless of its position in the trough. If cathodal shocks could cause accumulation of unknown substances in the cathodal half of the trough, this control group should show the same behavior as the experimental group. Interestingly, this control group showed the same behavior as the no-shock control group. Therefore, this possibility seems to be ruled out.

Second, it is possible that the mere presence of paramecium in one side of a trough can cause accumulation of its metabolites (e.g. carbon dioxide) which will attract the organism to one side of the trough (due to a decrease in PH levels which is attractive for paramecium since it can be a sign of bacterial food source (11)). To address this issue, Armus et al. have shown that changing the bright and dark side of the trough in test trials does not alter the tendency of paramecium to spend time on the bright side of the trough (Figure 6).

**Figure 6.**
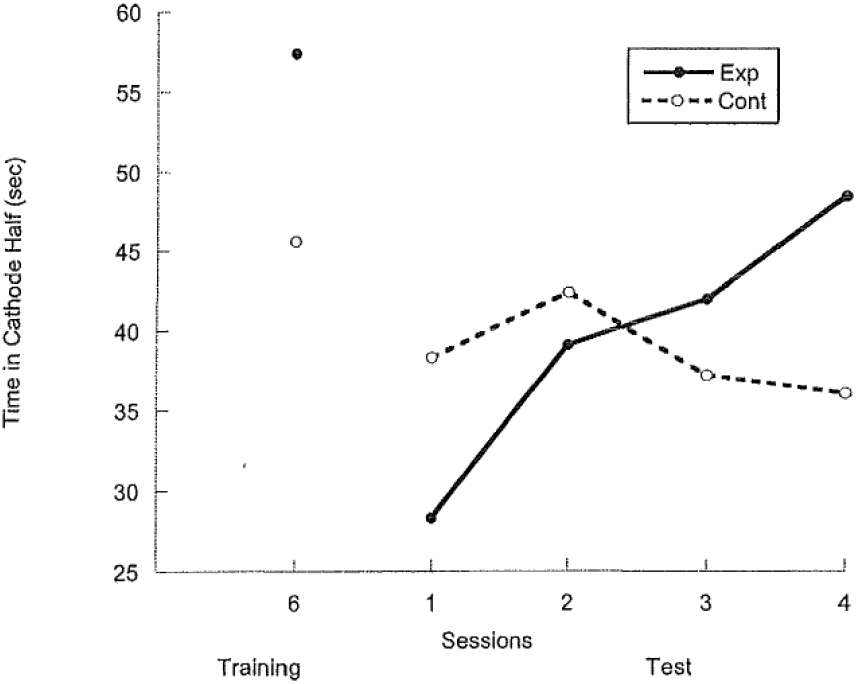
the mean time spent in the cathodal half of the trough in 4 test trials (and the last training trial) when the place of the cathodal half changed for test trials (Dashed line is the control group). Data suggest that the paramecium still shows a “tendency” to spend time in the cathodal half of the trough even though the location of the cathodal half was changed. adapted with permission from [6].

### Photoreception in P. caudatum

Light sensitivity in paramecium was reported by Jenings more than a hundred years ago (11). Armus et al. report (6) and its replication (12) are just some recent reports of this capability. It is known that light exposure can induce or modulate biological processes in cellular structures that do not possess a structurally distinct light detection system. This includes growth stimulation in yeast cells(13), activation of pig’s neutrophils(14), and growth modulation in paramecium itself (15). There is also a molecular model to explain this phenomenon (16). Accordingly, we suggest that a photoreception system may exist in paramecium. However, molecular pathways of such a system are still unknown. We believe that exploration of the evolutionary relationship between “photoreceptive unicellular organisms” can help here.

As described in (17), there is an eyespot apparatus (EA) called “stigma” in some of the motile photosynthetic organisms and green algae. The eyespot apparatus mediates the phototactic movements of the organism through molecular cascades. For instance, it is reported that in flagellated alga Chlamydomonas reinhardtii, light activates a signaling cascade involving archaeal-type rhodopsin (18). In euglena (a unicellular photosynthetic organism), it has been shown that light avoidance is mediated through a blue-light-activated adenylyl cyclase and cAMP (19). This blue-light receptor flavoprotein is the light receptor in euglena. Accordingly, it seems that the cAMP is an integral part of photo orientation processes in several unicellular organisms.

Another important signaling agent in phototaxis process seems to be the Ca^2+^ ion which is assumed to be one the major signaling mediators in plants and animals (20–22). Ca^2+^ is believed to be involved in the light modulated movement of green algae (particularly in Chlamydomonas) (23–25).

It is remarkable that eukaryotes have achieved the capability of phototaxis independently for at least eight times (17) and it is not difficult to achieve such a capability(17). In Ciliates, the phototactic activity can depend on the nutritional status of the organism in a way that the under-fed organism forms stigma and symbiotic relationship with a green alga and shows phototaxis towards the light source. On the other hand, well-fed organisms digest the stigma, hold the photoreceptors and exert a negative phototaxis. This probably helps the organism to feed its symbiont during under-fed situations and lose it in well-fed situations. Interestingly, *Paramecium bursaria* forms a similar symbiotic relationship with the green alga Zoochlorella i.e. when the environment suits photosynthesis, P. bursaria forms a symbiotic relationship with the Zoochlorella and when environmental parameters are not suitable for photosynthesis, P. bursaria digests its symbiont. The mechanism of steering in ciliates is still unknown but it has been suggested that there are light sensing vesicles that form an independent miniature stigma with their associated cilia (17).

Based on the aforementioned lines of evidence, we argue that paramecium possesses a similar light detection system that includes an unknown photoreceptor molecule and cAMP. In the following, we will suggest a molecular cascade based on our data to explain the light detection and learning capability in P. caudatum.

### Molecular pathways of learning in paramecium

Freely swimming paramecium spend around 39% of a trial’s time on the bright side of the trough (based on our data). Therefore, it might be possible to assume that paramecium has a “photophobic behavior”. Accordingly, we suggest that light exposure might increase cAMP concentration and since cAMP will increase the ciliary beat frequency in paramecium (26), light exposure can potentially increase paramecium’s swimming speed in the bright areas. This causes the paramecium to leave the bright side of the trough faster than its dark side which causes an overall reduction of time spent in the bright side of the trough. Moreover, another major role player in paramecium’s movement is voltage-gated Ca^2+^ channels (27, 28). It is known that membrane depolarization causes a reversal in ciliary beating direction of paramecium through Ca^2+^ (29). Since the resting membrane potential of the paramecium is around −25 millivolts (26), cathodal shocks can depolarize paramecium’s membrane. Therefore, cathodal shocks can reduce paramecium’s swimming speed through abovementioned mechanisms and cancel the “light-induced speed increase” (see Figure 7)

**Figure 7.**
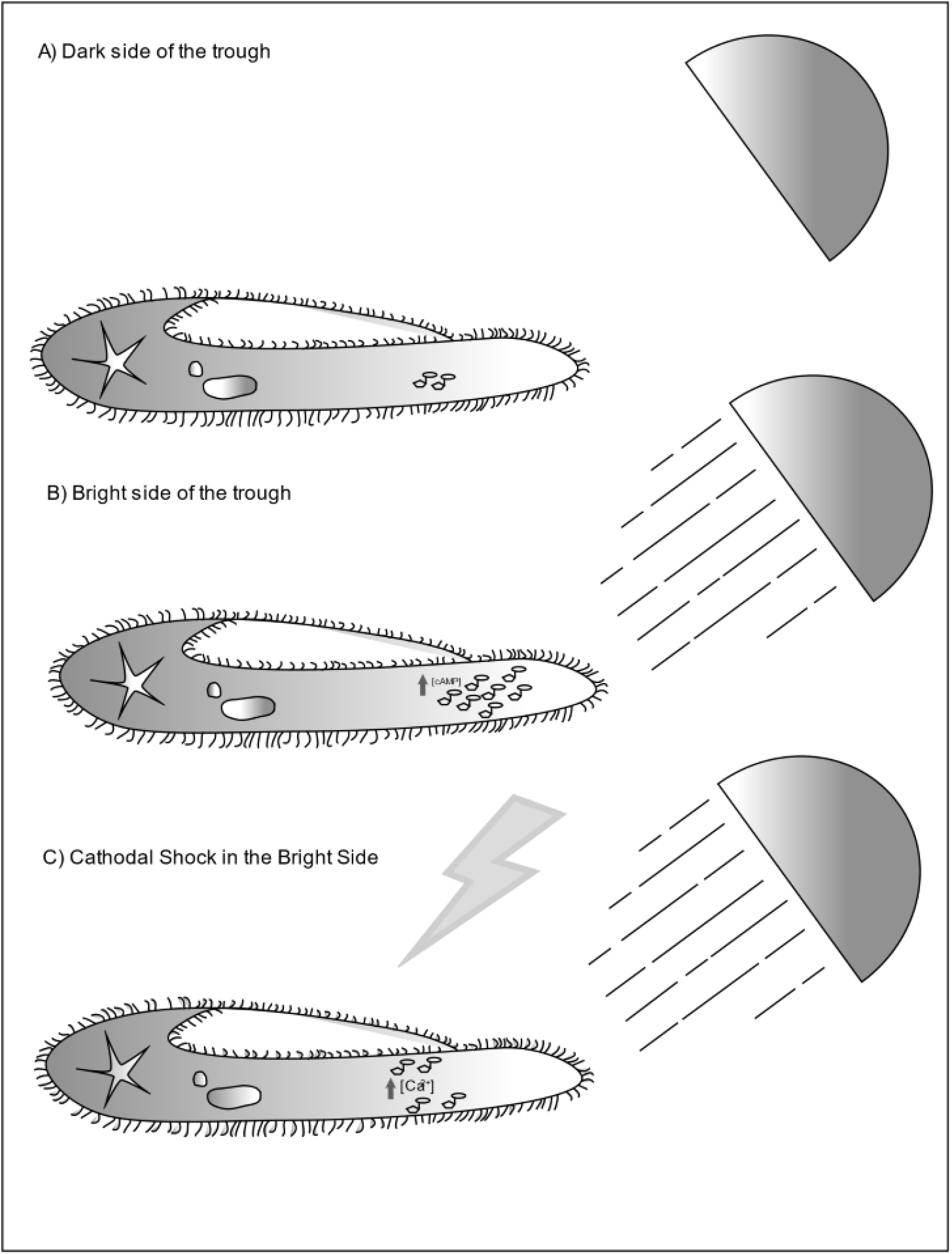
Proposed learning mechanism in paramecium. A: swimming in a relatively dark area maintains a minimal cAMP concentration. B: when paramecium enters the bright side of the trough, light exposure causes an increase in intracellular cAMP levels and increased swimming speed. C: If paramecium receives cathodal shocks when it is swimming in the bright side of the trough, the electrical shocks will cause subtle and temporary backward movements through miniature depolarization of the membrane and calcium inward flow. This will cancel out the increased swimming speed in bright side of the trough. Additionally, this process causes accumulation of cAMP in intracellular environment which leads to cancellation of photophobic behavior in paramecium during test trials.

However, this does not explain the capability of paramecium to retain information after the training trials. How does paramecium keep the learned information? We propose that when paramecium spends more time on the bright side of the trough, more cAMP can accumulate in the cytosol due to light exposure. Therefore, during the test trials, accumulated cAMP molecules will act as *memory molecules* and boost the swimming speed of the paramecium regardless of its position in the trough. This is in line with the experimental finding that paramecia in experimental group spend an almost equal amount of time in both halves of the trough during the test trials (56%, see the results section and Figure 3 for more detail).

According to our hypothesis, since cathodal shocks are countering the assumed cAMP-driven photophobic behavior in the bright side of the trough, paramecium cannot learn to associate the dark side of the trough with cathodal shocks. This prediction was tested and validated in the dark association group experiment (Figure 4).

## Conclusion

In conclusion, our results in line with other studies suggest that paramecium can have associative learning. Meanwhile, there are several questions about learning in paramecium that need to be addressed i.e what is the exact molecular pathway that governs this behavior? Is there a similarity between learning mechanisms in paramecium and other animals that could potentially translate to Alzheimer’s disease research? These questions can be addressed through pharmacological manipulations of paramecium learning. Future studies on the mechanisms of paramecium learning can shed more light on our understanding of learning at the molecular level in unicellular organisms.

